# What drives results in Bayesian morphological clock analyses?

**DOI:** 10.1101/219048

**Authors:** Caroline Parins-Fukuchi, Joseph W. Brown

## Abstract

Recently, approaches that estimate species divergence times using fossil taxa and models of morphological evolution have exploded in popularity. These methods incorporate diverse biological and geological information to inform posterior reconstructions, and have been applied to several high-profile clades to positive effect. However, there are important examples where morphological data are misleading, resulting in unrealistic age estimates. While several studies have demonstrated that these approaches can be robust and internally consistent, the causes and limitations of these patterns remain unclear. In this study, we dissect signal in Bayesian dating analyses of three mammalian clades. For two of the three examples, we find that morphological characters provide little information regarding divergence times as compared to geological range information, with posterior estimates largely recapitulating those recovered under the prior. However, in the cetacean dataset, we find that morphological data do appreciably inform posterior divergence time estimates. We supplement these empirical analyses with a set of simulations designed to explore the efficiency and limitations of binary and 3-state character data in reconstructing node ages. Our results demonstrate areas of both strength and weakness for morphological clock analyses, and help to outline conditions under which they perform best and, conversely, when they should be eschewed in favour of purely geological approaches.

## Introduction

Divergence time studies that incorporate morphology and more extensive fossil information have exploded over the last decade. These have sought to improve the dramatic gap between molecular estimates of divergence times and the fossil record. By integrating combinations of fossil preservation, lineage diversification, and morphological models, these studies have yielded a deeper understanding of the capability of fossil data to inform phylogenetic relationships, the timing of major radiations, and evolutionary patterns between fossil and living taxa [1-6].

In these approaches, discrete morphological data are analysed by calibrating substitution rates calculated under Markov substitution models and Poisson clock models to infer divergence times [1,2]. Models employed in these approaches may either assume a ‘strict’ clock, where rates remain constant across all lineages, or a ‘relaxed’ clock, where rates are allowed to across branches [7]. In current implementations, morphological clocks participate along with Bayesian tree priors to reconstruct posterior divergence times. The tree priors most commonly used are variations of ‘birth-death serial sampling’ (BDSS) models. These incorporate diversification and fossil sampling processes and often accommodate sequential sampling of ancestral taxa [3,5,8,9]. The more thorough integration of geological information enabled through these priors, which may be represented by temporal occurrence ranges or point appearances, seems to increase the accuracy and internal consistency over previous morphological dating methods, and has contributed to a more complete understanding of the evolution of several major clades [5,6]. Though total-evidence dating methods were originally developed to analyse fossil taxa alongside living taxa, they have been increasingly applied to exclusively palaeontological datasets, where they are used to infer divergence times from morphological data alone [10]. Palaeontologists also frequently employ *a posteriori* time-scaling (APT) approaches that scale cladograms to time. These scale branches either directly to observed ranges [11,12] or model diversification and fossil preservation to estimate divergence times probabilistically [13]. The latter approach is a variant of the tree priors described above but differs by its exclusion of a morphological clock.

Despite the recent surge in popularity experienced by the many permutations of morphological clock approaches, there remain outstanding challenges in their use. Some of these are shared with molecular data, and others are unique to morphology. Complex interactions between fossil calibrations and other priors can create conflict, yielding results which are egregiously incorrect yet overly precise [14]. In a similar vein, the accuracy and precision of molecular and morphological clock estimates can be driven by prior choice and the reliability of the fossil record [15,16]. This raises questions about the degree to which character data contribute to posterior divergence time estimates relative to prior model choice and temporal information gleaned directly from fossils. This is especially important for morphological characters, for which complexities in evolutionary processes, errors stemming from character coding, and biases in sampling can create a mismatch between data and simple evolutionary models, potentially leading to unrealistic divergence time estimates [4]. These difficulties can result in imprecise, unrealistically ancient estimates that are only improved when constrained by strongly informative priors [17].

Results achieved from BDSS tip-dating can differ substantially from APT methods [18]. These differences are unsettling given the lack of a theoretical basis from which to expect clock-like evolutionary patterns in morphology. While molecular dating can sometimes yield muddled results, these methods possess a strong theoretical grounding [19,20]. On the other hand, some datasets reveal striking congruence between geology and dates derived under BDSS methods [10]. These observations beg question as to the conditions under which morphological clock approaches provide substantial information when combined with geological dating methods.

In this study, we examine the capability of morphological characters to inform divergence times. We examine this issue in a Bayesian context by dissecting the relative contribution of morphological clocks and prior information derived from complex models of speciation and fossil preservation to posterior divergence time estimates. In Bayesian analyses, characterising the extent to which prior beliefs contribute to posterior estimates can help isolate the contribution of information presented by new data [14,21]. We perform these comparisons using three morphological datasets representing canids, cetaceans, and hominins to compare posterior estimates to those estimated under geologically-informed priors alone. We supplement these examinations with a set of simulations designed to gauge the amount of character data needed to inform the ages of internal nodes in the absence of sampling bias and model misspecification.

## Materials and Methods

### Empirical datasets

We obtained three empirical datasets from the literature containing morphological and geological range data in canids [10,22], hominins [23], and cetaceans [24].

### Divergence time estimation

We estimated divergence times and topology using BEAST 2 (version 2.4.7) [25] (Fig. 1). Topology was initially estimated using the full character and stratigraphic range datasets. In subsequent analyses that estimate ages using reduced and simulated datasets, topology was constrained to the initial result to make node age comparisons more straightforward. Tip dates were indicated as the most recent occurrence of each taxon in the fossil record. Stratigraphic ranges were specified for all fossil taxa after [10], using first appearance dates (FAD) and last appearance dates (LAD) that were published along with the morphological matrices for all datasets.

**Figure 1.**
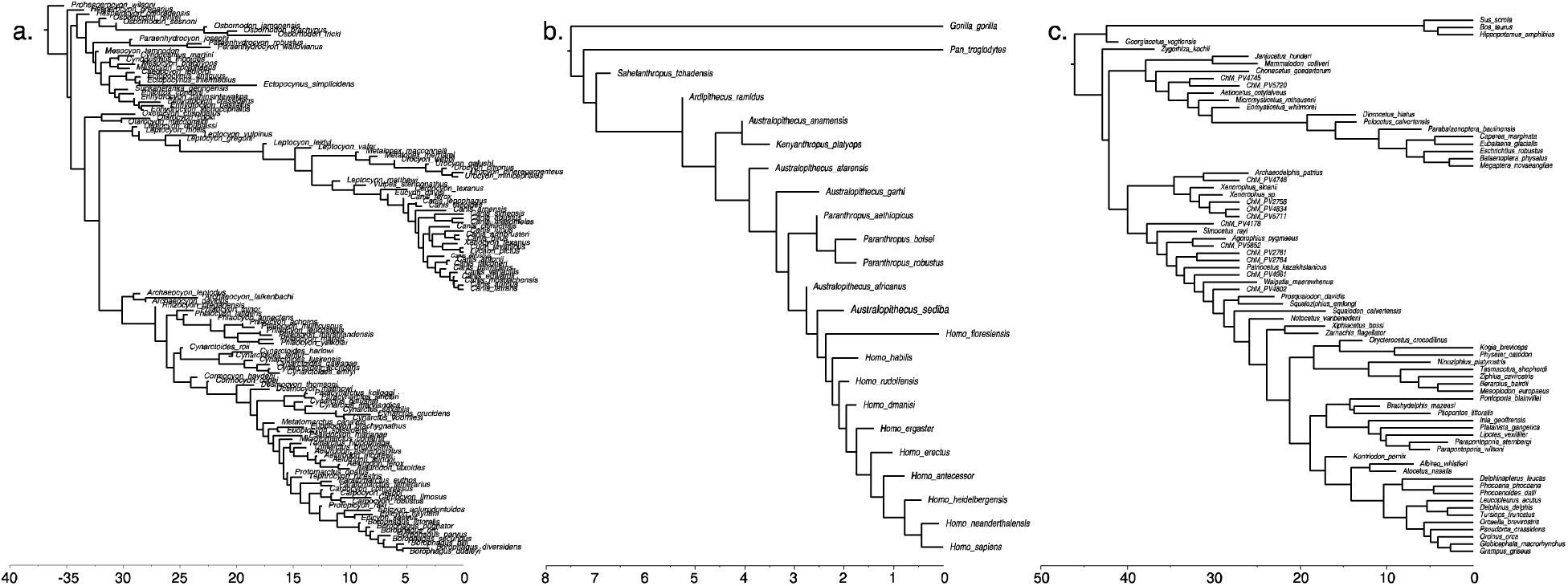
Maximum clade credibility posterior reconstructions of topology and divergence times in a) canids, b) hominins, and c) cetaceans. The Fossilized Birth-Death parameters were calibrated using stratigraphic ranges of the fossil tips.

For all three datasets, divergence times were estimated using an uncorrelated log-normal (UCLN) relaxed clock. We used the ‘Fossilized Birth-Death’ (FBD) prior deployed in the BEAST 2 ‘Sampled Ancestors’ package (version 1.1.7) [3,9]. Prior distributions for each parameter were chosen to be informative as follows. For the canid dataset, we retained the prior values used by [10] in their analysis. Diversification and extinction rate values were chosen in hominins to reflect the high extinction that is apparent in their fossil record. In the cetaceans, these priors were informed by previous diversification studies[26]. Example BEAST 2 control files are provided in the data supplement. Markov Chain Monte-Carlo (MCMC) simulations were run between 30 and 60 million generations. All runs were manually checked for convergence using Tracer v1.6 [27], and accepted when all logged parameters reached an effective sample size (ESS) of at least 200. To further examine the behaviour of mechanistic priors and morphological clocks on posterior ages, we uniformly increased the FAD of each temporal occurrence range used for the prior calculations in the cetacean dataset by 10 Ma. We then reconstructed ages using this altered geological information, following the same procedure as is described above.

### Simulation

To better explore the abilities and limitations of character data to inform divergence times in the absence of complicating factors such as model misspecification and sampling bias, we performed a small series of simulations. Using the tree estimated from the empirical cetacean matrix, we artificially increased the height of two nodes, one close to the present, and one nested deeper within the tree (Fig. S1). This modified tree was used to simulate matrices of 1000 characters. These characters were randomly subsampled into matrices of 10, 30, 70, and 500 characters to examine the ability of character data to inform posterior ages at varying levels of abundance. To examine the effect of differing discrete character state space, we performed this test using binary and 3-state characters. Characters were all simulated under the Mk model of morphological evolution [28] using the ‘geiger’ R package [29] with clock-like rates across the tree. Matrices were generated using two different schemes of site-wise rate variability. One set was evolved along a single rate across all traits, and another using five separate rates. The multi-rate matrices were generated by concatenating five 200-character matrices evolved using distinct rates, and were randomly subsampled to yield the smaller datasets. R and Python scripts used for simulation are available in the data supplement. To ensure the informativeness of the simulated characters, trees were inferred from all matrices using RAxML version 8.2.11 [30] and visually checked for topological and branch length accuracy. We then inferred divergence times from the simulated matrices using BEAST 2, following a similar procedure as for the empirical datasets. However, since the characters were simulated to be clock-like, we inferred dates under a strict clock rather than a UCLN relaxed clock to avoid imprecision stemming from over-parameterization. All priors were the same as those for the empirical analyses. This enabled testing of the required number of characters needed to reconstruct the single altered internal node age when sampling is complete, and modelling assumptions are not violated.

## Results and Discussion

### How do morphological clocks drive posterior age estimates?

For the canid and hominin datasets, we find that posterior divergence times are not substantially different from those estimated from the prior model alone (Fig. 2). This is the case when analysed using priors both with (Fig. 2a) and without (Fig. 2b) temporal ranges. There exist a small handful of deviations from this pattern, but nearly all residuals fall within 1 Ma of the identity line. 95% credibility intervals (CI) are wider when temporal ranges are included in the prior. Precision in both tip and internal node ages is slightly higher in posterior estimates compared to posterior. The weak influence of morphological data on posterior divergence time estimates may provide an explanation for the strong internal consistency found by [10] through cross validation (CV). In that study, the authors attribute their results to strong performance of the morphological clock. In their procedure, the authors estimated the age of fossils when geological information is removed from a single tip. Based upon our results, their procedure might be interpreted as testing the consistency of the FBD prior, rather than the morphological clock. The hominin matrix differs in that, when stratigraphic ranges are incorporated into the prior, morphological data push dates slightly older for nodes close to the present, but the disparity between prior and posterior estimates decreases toward the root (Fig. 2c). In the absence of stratigraphic information, posterior and prior mean ages align closely with one another (Fig. 2d). Thus, the extension of geological point occurrences to ranges appears to increase the amount of information extracted from morphological characters.

**Figure 2.**
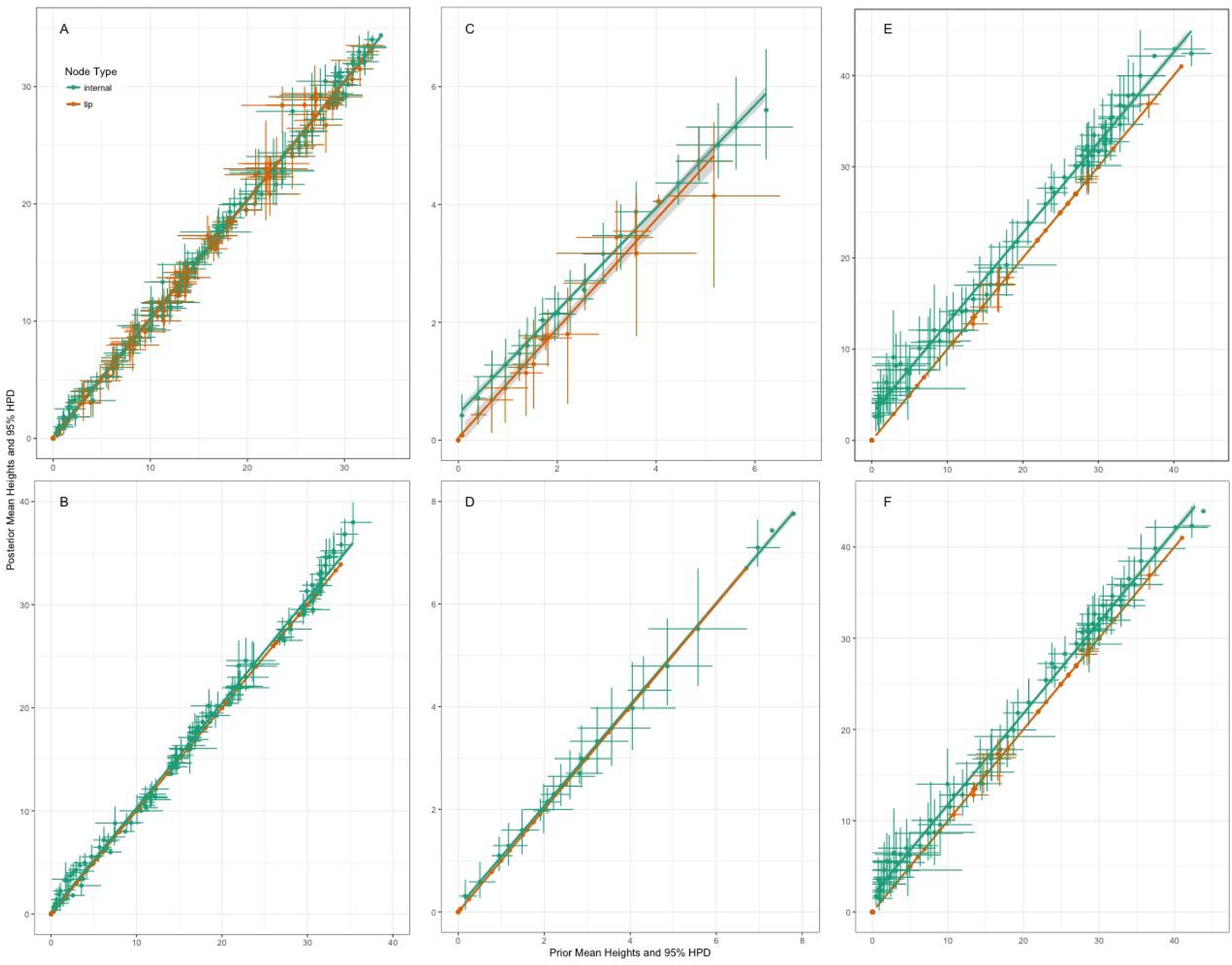
A,B) Canid prior and posterior node ages. Node heights reconstructed with A) full stratigraphic ranges for fossil tips and B) without full stratigraphic ranges. When ranges were not used, tip calibrations were added as the LAD of the ranges. C,D) hominin prior and posterior node ages reconstructed C) with and D) without stratigraphic ranges. E,F) cetacean prior and posterior node ages reconstructed E) with and F) without stratigraphic ranges.

In contrast to the hominid and canid datasets, the cetacean morphological data pushed divergence times older than prior estimates. Prior mean node ages that incorporate temporal ranges (Fig. 2e) were older than those estimated without ranges (Fig. 2f). For both range-informed and range-uninformed priors, addition of morphological data increased mean ages by approximately 0-3 Ma across the tree. When the upper range of the geological ranges were altered to be 10 Ma older across all taxa, the posterior followed the prior. Posteriors estimated under the altered priors exhibited a similar pattern to the analyses with unaltered priors, with posterior and prior ages differing by approximately 0-3 Ma across all nodes (Fig. S2). Since posterior estimates shifted in tandem with changes in priors, the relaxed morphological clock seemed to inform relative, rather than absolute, divergence times. Absolute divergence times were then made identifiable by geologically-informed priors.

### Simulations

We found that, overall, binary characters were less informative in divergence time reconstruction than trinary characters (Fig. 3). At nearly every matrix size and rate configuration, ages estimated from the 3-state characters were older and closer to the true node age compared to the binary characters. This was the case for both the shallow and deep nodes. The fewer number of states present in binary characters appears to result in decreased information. This is expected from Shannon information theory [31] where information content limits are determined by the size of the alphabet involved. Biologically, this result may be due to the greater propensity toward rapid saturation exhibited by binary characters, making it difficult to correctly estimate the number of changes undergone at each site. This suggests that, all things being equal, researchers should be cautious when interpreting divergence times estimated from matrices which are primarily composed of binary characters, as they have a greater propensity to underestimate true divergence times compared to characters with larger numbers of states. However, the binary portion of the empirical cetacean matrix appears to retain dating information. This may illustrate that the complexities in morphological evolution and character sampling may yield unpredictable results, complicating the ability to make general prescriptive statements. Nevertheless, combined with the empirical results described above, the weak patterns observed in the simulations further underscore the importance in comparing priors and posteriors.

**Figure 3.**
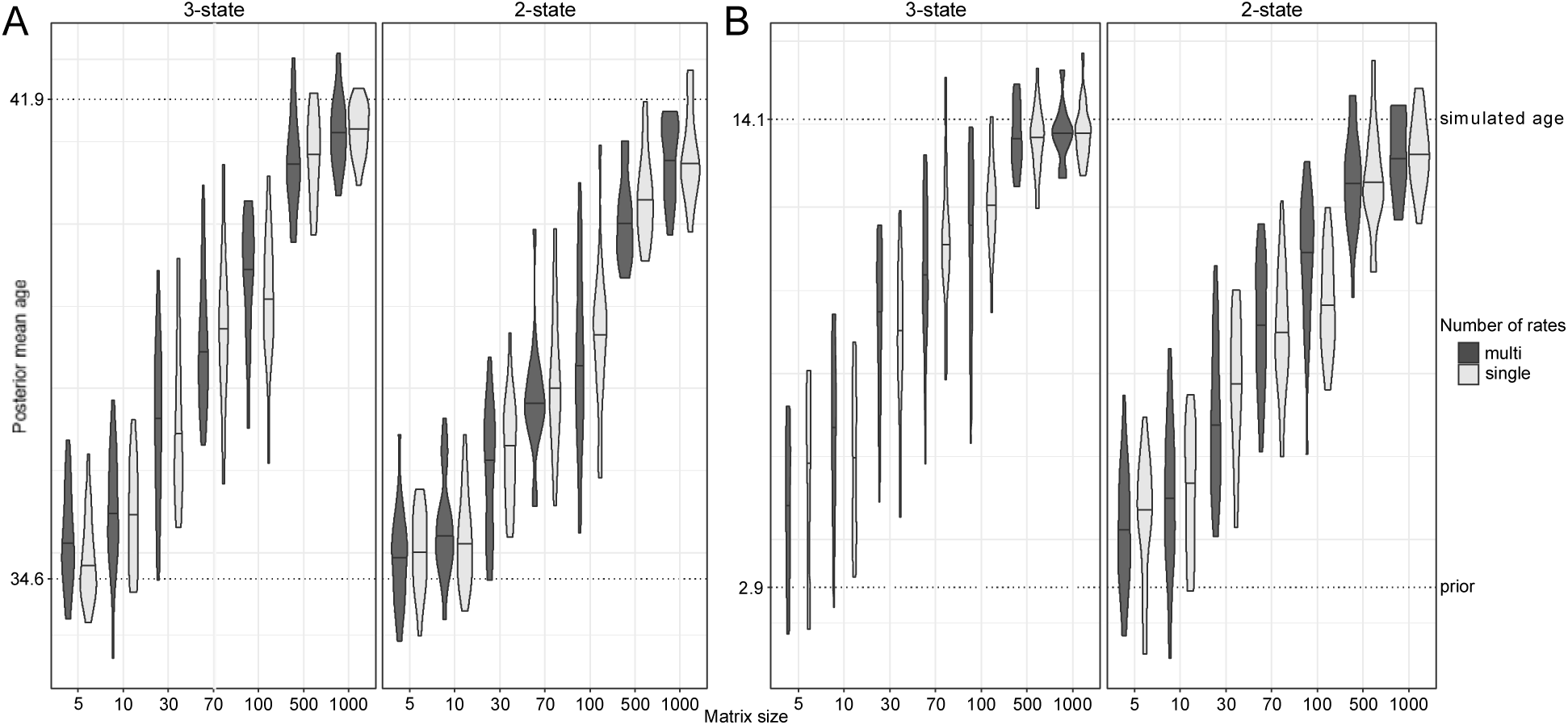
Comparison between “true” simulated ages, and mean ages reconstructed from simulated data. A) reconstructions for the deep node shown in figure 2. B) shallow node reconstructions. Plots are subdivided by the state space of the data matrix. Violin plots are coloured by number of rates simulated within each dataset. Comparisons were performed between datasets simulated under a single rate, and under multiple randomly selected rate categories. Dotted lines indicate the ages the data were simulated under (upper) and ages recovered under the prior alone (lower).

### Homoplasy, missing data, and information content

Information content varies across these three matrices. While the cetacean matrix is intermediate in size, as compared to the other two, it differs in sampling completeness, statistical behaviour, and the distribution of substitutions across characters. These properties may have resulted in the increased information in the cetacean morphological matrix. The three datasets differ substantially in their respective proportions of missing data. The hominin matrix is the sparsest, with over half of sites missing for each taxon on average (Fig. 4a). Gappiness is distributed widely, ranging from ∼20% to nearly all sites missing. The canid distribution possesses the lowest proportion of missing sites. The cetacean dataset is further distinguished from the hominin dataset in the tameness of its statistical behaviour. Relaxed clock parameter estimates show that the cetacean data were relatively clock-like compared to the other datasets (Table 1). The hominin matrix possesses much greater branch-wise rate heterogeneity, possessing nearly three times greater variability across branches in morphological rate compared to cetaceans. It is possible that this deviation from the morphological clock limits the amount of recoverable information about divergence times.

**Figure 4.**
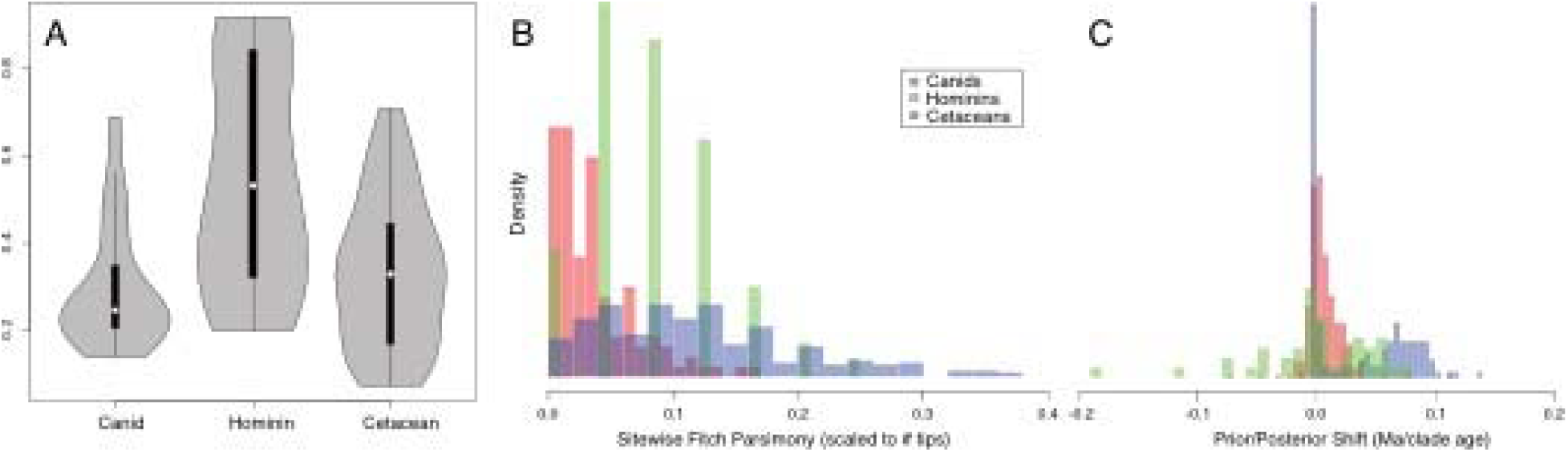
A) Proportion of missing characters for each taxon across each dataset. B) Distribution of number of parsimony changes across sites for each dataset. These values are scaled to the number of tips represented in each matrix. C) Distribution of shift between prior and posterior mean node age estimates. Values were calculated by subtracting posterior from prior heights and scaling the values to the overall depth of the tree. Histograms represent the density rather than raw frequency counts.

**Table 1.** Mean rates and variance estimated under the Uncorrelated Lognormal (UCLN) model.

| Dataset | Mean (Ucld.mean) | Variance (Ucld.stdev) |
| --- | --- | --- |
| Canids | 0.09426 | 1.163 |
| Cetaceans | 0.04184 | 0.509 |
| Hominins | 2.125 | 1.56 |

The cetacean matrix contains greater variability in the number of character changes across all characters (Fig. 4b). The comparatively large number of substitutions implied by the cetacean matrix may increase the chance of recovering a character change along any single branch, increasing the amount of information from which to condition the clock model. The canid and hominin datasets, on the other hand, have low homoplasy relative to the Bayesian summary topology, a pattern that may be common to many cladistic morphological datasets. These patterns demonstrate opposing challenges in applying clock models to the analysis of existing morphological datasets. Although the canid and hominin data show patterns that suggest high *topological* information content, the conserved number of character changes limits the capability to inform divergence times. On the other hand, the higher entropy displayed by the cetacean matrix would be expected to provide clock models with more information (Fig. 4c). Thus, if information content is conceived using intersecting axes of character change count and completeness of character sampling, the cetacean dataset possesses the largest amount of information among these three datasets.

However, it is difficult to determine whether this added information is likely to result in increased accuracy in divergence time reconstructions. There is no empirical or theoretical expectation that morphological characters evolve concordantly to even relaxed clocks. Instead, certain lineages can be intensely biased toward rapid and major morphological rearrangement, while others may remain morphologically static [32]. As a result, the increased variation in pattern displayed by the cetacean dataset may reflect either accurately informative signal, random noise, or procedural bias in evolutionary change. Similarly, it is possible for even random data to inform divergence times, albeit inaccurately. These complications may be the fundamental source of the strongly misleading results generated during previous authors’ attempts to apply morphological clocks to date the mammalian radiation [4,17]. The matrix used in these previous studies compiled a massive number of characters from widely different sources, likely resulting in extremely high entropy. Complex patterns and noise in these data may require unrealistic ancient dates in order to fit under a clock model. This interpretation is underscored by the wide behavioural disparity observed in each of our analyses. It is difficult to envision that existing methods would be capable of recovering accurately informative signal from a hypothetical concatenated analysis of these three datasets. These problems highlight the difficulty in interpreting the accuracy of morphological clocks in reconstructing dates, even in apparently well-behaved datasets such as the cetaceans.

### Rocks vs morphological clocks

The results presented here demonstrate that geological data provide the most reliably accurate information for divergence time estimation. In the canid and hominin dataset, we find that morphological data are limited in their ability to inform divergence times, and so geological occurrences are a more reliable and consistent data source. The cetacean matrix more informs posterior divergence time estimates more substantially. However, our comparison of the results achieved under different priors demonstrates the crucial role of geological data in cases where accuracy and precision are needed when estimating absolute divergence times from morphological data. This is true even in cases where morphological data appear to be well sampled and behaved, such as is shown in the cetacean dataset.

In contrast to the geological data, the morphological matrices examined here expose limitations. The weakly informative canid and hominin morphology raises questions concerning the trade-offs in incorporating morphological data in divergence time estimation. Our study highlights several risks in the use of morphological clocks. In cases such as our canid analysis, where morphological data appear to behave consistently but are uninformative, analysis of morphological clocks in a Bayesian context may yield misleadingly high confidence that posterior dates reflect true speciation times. Although the simulations remove model misspecification and incomplete character sampling as a potential source of error, a relatively large number of characters are needed to substantially inform posterior estimates. Researchers using these approaches should take extra caution to compare prior and posterior estimates to determine whether posterior signal originates from the data or the prior, a concern that involves molecular data as well [14]. Furthermore, difficulty in distinguishing between evolutionary signal and noise and lack of a theoretical expectation remains a source of substantial and fundamental underlying concern, even in cases where data seem internally consistent and well-behaved.

Our simulations yielded the insight that morphological clock methods to underestimate true divergence times, even at large matrix sizes. Since many morphological matrices contain only small numbers of characters, many datasets may be incapable of recovering true divergence times, even when egregious sampling biases and model violations are not present. Thus, dates recovered using from morphological clocks using even well-behaved empirical datasets should be interpreted cautiously. For example, although the empirical cetacean character matrix analysed here consistently results in date estimates that are several million years older than those recovered under the prior, it is possible that these dates still represent underestimates of true divergence times, and should therefore still be conservatively interpreted as minimum divergence times, similarly to typical treatments of geological estimates. This may also extend to previous studies which infer recent dates for radiations using total-evidence methods [6].

### How can we move forward?

Our analyses demonstrate several challenges in recovering divergence times that are unique to morphological data. Many existing morphological datasets, including the canid and hominin matrices analysed here, are cladistically informative, but lack clock information. Other datasets may be more informative, but the lack of a theoretical expectation makes their interpretation unsettling. When reconstructing divergence for older clades than examined here, these problems may occur simultaneously, yielding unpredictable results. We recommend that researchers approach morphological clock estimates with caution, and estimate dates from geological information alone when in doubt. The poor trade-offs demonstrated by morphological clock methods and dilemmas in the interpretation of their results suggests that their use should remain limited. New developments will continue the existing synthesis of neo- and palaeontological data within the framework encapsulated by existing approaches while generating unprecedented discoveries.

## Acknowledgements

CPF and JWB would like to thank JF Walker and SA Smith for helpful comments that improved the manuscript.

## Data, code, and materials

Scripts used for simulation, data processing, and statistical analyses can be accessed at https://github.com/carolinetomo/mammalian_morphological_clocks. Example BEAST 2 control files are available on Dryad.

## Author contributions

CPF conceived of and designed the study, carried out dating analyses and simulation experiments, performed statistical analyses, and drafted the manuscript. JWB contributed to dating analyses, interpretation of results, and helped draft the manuscript.

## Supplementary Figures

**Figure S1.**
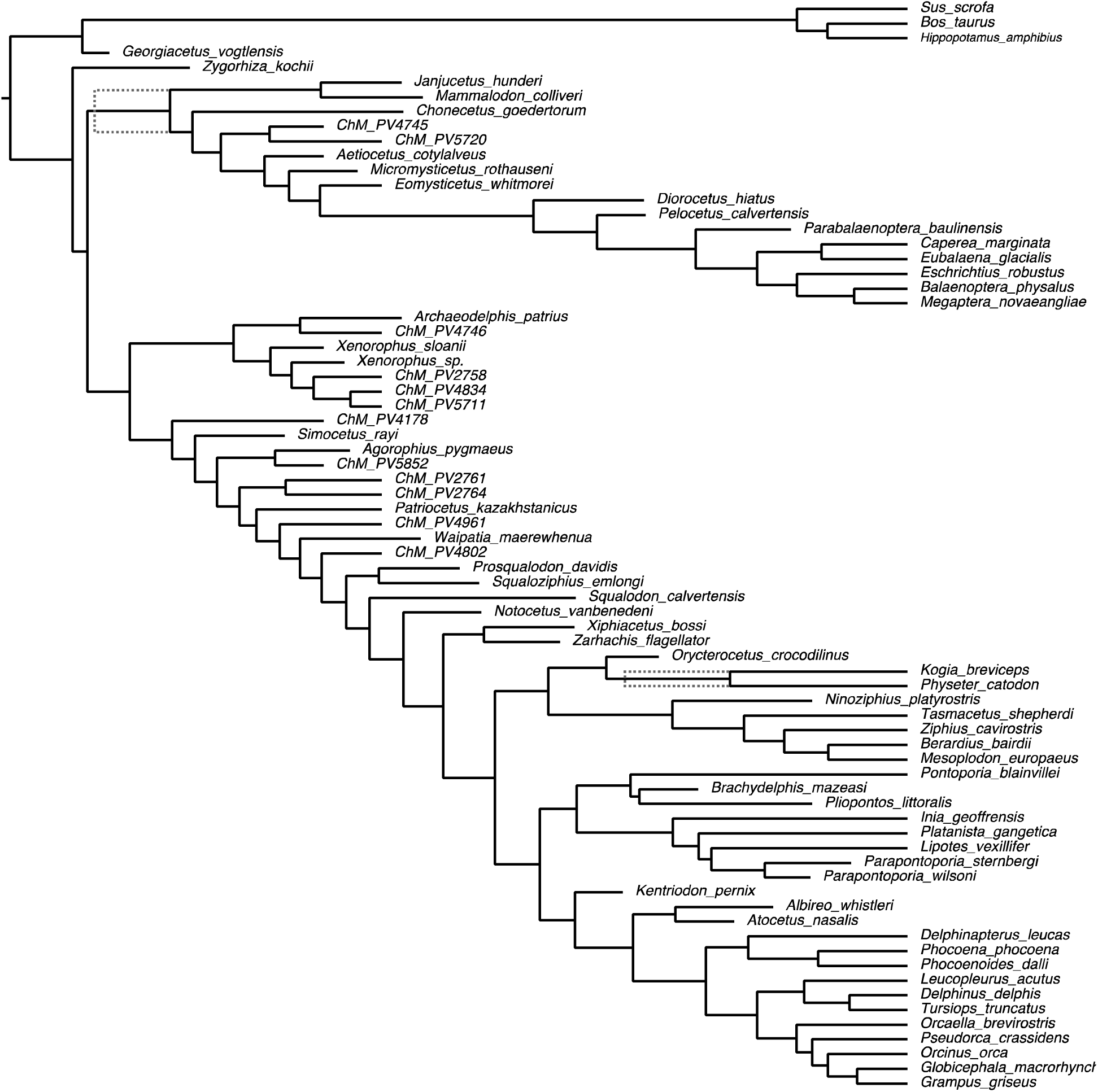
Nodes altered for simulations. Grey dotted lines show altered node heights.

**Figure S2.**
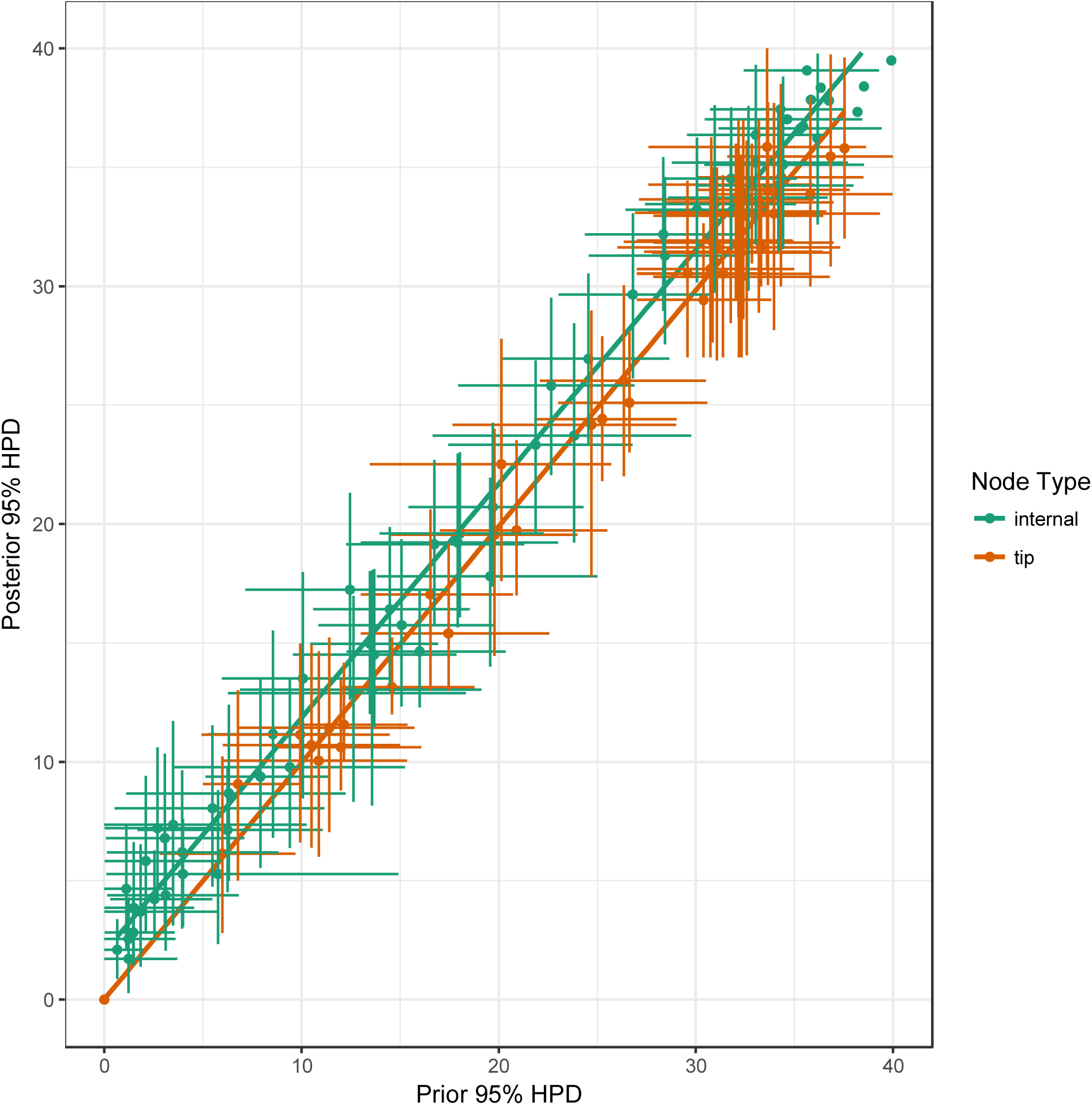
Prior and posterior mean age reconstructions with all FADs altered to be 10 Ma older.

## References

1. Pyron, R. A. 2011 Divergence time estimation using fossils as terminal taxa and the origins of Lissamphibia. Syst. Biol. 60, 466–481. (doi:10.1093/sysbio/syr047)

2. Ronquist, F., Klopfstein, S., Vilhelmsen, L., Schulmeister, S., Murray, D. L. & Rasnitsyn, A. P. 2012 A total-evidence approach to dating with fossils, applied to the early radiation of the hymenoptera. Syst. Biol. 61, 973–999. (doi:10.1093/sysbio/sys058)

3. Heath, T. A., Huelsenbeck, J. P. & Stadler, T. 2014 The fossilized birth-death process for coherent calibration of divergence-time estimates. Proc. Natl. Acad. Sci. U.S.A. 111, E2957–66. (doi:10.1073/pnas.1319091111)

4. Beck, R. M. D. & Lee, M. S. Y. 2014 Ancient dates or accelerated rates? Morphological clocks and the antiquity of placental mammals. Proc. Roy. Soc. B 281, 20141278–20141278. (doi:10.1098/rspb.2014.1278)

5. Zhang, C., Stadler, T., Klopfstein, S., Heath, T. A. & Ronquist, F. 2016 Total-Evidence Dating under the Fossilized Birth-Death Process. Syst. Biol. 65, 228–249. (doi:10.1093/sysbio/syv080)

6. Gavryushkina, A., Heath, T. A., Ksepka, D. T., Stadler, T., Welch, D. & Drummond, A. J. 2017 Bayesian Total-Evidence Dating Reveals the Recent Crown Radiation of Penguins. Syst. Biol. 66, 57–73. (doi:10.1093/sysbio/syw060)

7. Drummond, A. J., Ho, S. Y. W., Phillips, M. J. & Rambaut, A. 2006 Relaxed Phylogenetics and Dating with Confidence. PLoS Biology 4, e88. (doi:10.1371/journal.pbio.0040088)

8. Stadler, T. 2010 Sampling-through-time in birth–death trees. Journal of Theoretical Biology 267, 396–404. (doi:10.1016/j.jtbi.2010.09.010)

9. Gavryushkina, A., Welch, D., Stadler, T. & Drummond, A. J. 2014 Bayesian inference of sampled ancestor trees for epidemiology and fossil calibration. PLoS Comput. Biol. 10, e1003919. (doi:10.1371/journal.pcbi.1003919)

10. Drummond, A. J. & Stadler, T. 2016 Bayesian phylogenetic estimation of fossil ages. Philos. Trans. R. Soc. Lond., B, Biol. Sci. 371, 20150129. (doi:10.1098/rstb.2015.0129)

11. Clyde, W. C. & Fisher, D. C. 1997 Comparing the fit of stratigraphic and morphologic data in phylogenetic analysis. Paleobiology 23, 1–19. (doi:10.1017/s0094837300016614)

12. Huelsenbeck, J. P. & Rannala, B. 1997 Maximum Likelihood Estimation of Phylogeny Using Stratigraphic Data. Paleobiology 23, 174–180. (doi:10.2307/2401051?ref=search-gateway:0a54a52430893d9e4bc5fd3ff569737a)

13. Bapst, D. W. 2013 A stochastic rate-calibrated method for time-scaling phylogenies of fossil taxa. Methods in Ecology and Evolution 4, 724–733. (doi:10.1111/2041-210X.12081)

14. Brown, J. W. & Smith, S. A. 2017 The Past Sure Is Tense: On Interpreting Phylogenetic Divergence Time Estimates. Syst. Biol. (doi:10.1093/sysbio/syx074)

15. Condamine, F. L., Nagalingum, N. S., Marshall, C. R. & Morlon, H. 2015 Origin and diversification of living cycads: a cautionary tale on the impact of the branching process prior in Bayesian molecular dating. BMC Evol. Biol. 15, 65. (doi:10.1186/s12862-015-0347-8)

16. Warnock, R. C. M., Yang, Z. & Donoghue, P. C. J. 2017 Testing the molecular clock using mechanistic models of fossil preservation and molecular evolution. Proc. Roy. Soc. B 284, 20170227. (doi:10.1098/rspb.2017.0227)

17. Ronquist, F., Lartillot, N. & Phillips, M. J. 2016 Closing the gap between rocks and clocks using total-evidence dating. Philos. Trans. R. Soc. Lond., B, Biol. Sci. 371, 20150136. (doi:10.1098/rstb.2015.0136)

18. Bapst, D. W., Wright, A. M., Matzke, N. J. & Lloyd, G. T. 2016 Topology, divergence dates, and macroevolutionary inferences vary between different tip-dating approaches applied to fossil theropods (Dinosauria). Biol. Lett. 12, 20160237. (doi:10.1098/rsbl.2016.0237)

19. Kimura, M. 1968 Evolutionary Rate at the Molecular Level. Nature 217, 624–626. (doi:10.1038/217624a0)

20. Kimura, M. & Ohta, T. 1974 On Some Principles Governing Molecular Evolution. Proc. Natl. Acad. Sci. U.S.A. 71, 2848–2852. (doi:10.1073/pnas.71.7.2848)

21. Lewis, P. O., Chen, M.-H., Kuo, L., Lewis, L. A., Fučíková, K., Neupane, S., Wang, Y.-B. & Shi, D. 2016 Estimating Bayesian Phylogenetic Information Content. Syst. Biol. 65, 1009–1023. (doi:10.1093/sysbio/syw042)

22. Slater, G. J. 2015 Iterative adaptive radiations of fossil canids show no evidence for diversity-dependent trait evolution. Proc. Natl. Acad. Sci. U.S.A. 112, 4897–4902. (doi:10.1073/pnas.1403666111)

23. Dembo, M., Matzke, N. J., Mooers, A. Ø. & Collard, M. 2015 Bayesian analysis of a morphological supermatrix sheds light on controversial fossil hominin relationships. Proc. Roy. Soc. B 282, 20150943. (doi:10.1098/rspb.2015.0943)

24. Geisler, J. H., McGowen, M. R., Yang, G. & Gatesy, J. 2011 A supermatrix analysis of genomic, morphological, and paleontological data from crown Cetacea. BMC Evol. Biol. 11, 112. (doi:10.1186/1471-2148-11-112)

25. Bouckaert, R., Heled, J., Kühnert, D., Vaughan, T., Wu, C.-H., Xie, D., Suchard, M. A., Rambaut, A. & Drummond, A. J. 2014 BEAST 2: a software platform for Bayesian evolutionary analysis. PLoS Comput. Biol. 10, e1003537. (doi:10.1371/journal.pcbi.1003537)

26. Slater, G. J., Price, S. A., Santini, F. & Alfaro, M. E. 2010 Diversity versus disparity and the radiation of modern cetaceans. Proc. Roy. Soc. B 277, 3097–3104. (doi:10.1098/rspb.2010.0408)

27. Rambaut, A., Suchard, M. A., Xie, D. & Drummond, A. J. In press. Tracer v1.6.

28. Lewis, P. O. 2001 A Likelihood Approach to Estimating Phylogeny from Discrete Morphological Character Data. Syst. Biol. 50, 913–925. (doi:10.1080/106351501753462876)

29. Pennell, M. W., Eastman, J. M., Slater, G. J., Brown, J. W., Uyeda, J. C., FitzJohn, R. G., Alfaro, M. E. & Harmon, L. J. 2014 geiger v2.0: an expanded suite of methods for fitting macroevolutionary models to phylogenetic trees. Bioinformatics 30, 2216–2218. (doi:10.1093/bioinformatics/btu181)

30. Stamatakis, A. 2014 RAxML version 8: a tool for phylogenetic analysis and post-analysis of large phylogenies. Bioinformatics 30, 1312–1313. (doi:10.1093/bioinformatics/btu033)

31. Shannon, C. E. 1948 A Mathematical Theory of Communication. Bell System Technical Journal 27, 623–656. (doi:10.1002/j.1538-7305.1948.tb00917.x)

32. Simpson, G. G. 1944 Tempo and Mode in Evolution. New York: Columbia University Press.

